# Asynchronous cortical recovery from sleep inertia: region- and frequency-specific dynamics following deep NREM sleep

**DOI:** 10.64898/2026.02.08.704730

**Authors:** Chengmi Hu, Daisuke Shimane, Kiyoshi Nakahara, Masaki Takeda

## Abstract

Sleep inertia (SI) is a transient state of impaired alertness and cognition after awakening, typically exacerbated by prior deep sleep. Despite its prevalence, the fine-grained temporal dynamics of brain activity during this transition remains poorly understood. We employed trial-by-trial behavioral assessment (20-second resolution) to map the recovery trajectories of 44 participants following awakenings from either shallow (N1/N2) or deep (N3) NREM sleep. We found that awakening from deep NREM sleep resulted in a significantly slower return to baseline working-memory performance. This behavioral deficit was mirrored by specific EEG signatures: frontal low-frequency activity (delta/theta) returned to baseline levels within minutes, whereas posterior high-frequency activity (alpha/beta) remained altered for a prolonged period. These findings reveal that the human brain reboots asynchronously following deep NREM sleep, with a “frontal-first, posterior-last” gradient that dictates the time course of behavioral restoration.

## Introduction

Sleep inertia (SI) refers to the transitional state between sleep and full wakefulness that occurs immediately after awakening, characterized by impaired alertness and reduced cognitive function (Tassi & Muzet, 2000; Vallat et al., 2019). Prior research has identified several factors that modulate the severity and duration of SI, including sleep deprivation, alcohol consumption, and the sleep stage—rapid eye movement (REM) or non-rapid eye movement (NREM) at varying depths—immediately preceding awakening (Tassi & Muzet, 2000). Notably, both human and rodent studies emphasize that the pre-awakening sleep stage is a critical determinant, as it reflects the brain’s physiological state at the moment of transition to wakefulness (Human: Hilditch & McHill, 2019; Occhionero et al., 2021; Tassi & Muzet, 2000; Rodent: Vyazovskiy et al., 2014).

To date, the effects of SI have primarily been examined through behavioral performance and neural activity. Most studies have collected data at discrete intervals of several minutes immediately after awakening and quantified the magnitude of SI by comparing post-awakening measures to a pre-sleep baseline. Regarding behavioral outcomes, previous research indicates that SI impairs performance across a diverse range of cognitive tasks, with these deficits gradually dissipating over time (Bruck & Pisani, 1999; Groeger et al., 2011; Jewett et al., 1999; Liaukovich et al., 2022; Occhionero et al., 2021; Ritchie et al., 2017; Santhi et al., 2013; Tassi et al., 2006). Furthermore, SI tends to be more pronounced and persistent following awakenings from deep NREM sleep compared to shallow NREM sleep (Dinges et al., 1985; Hilditch & McHill, 2019; Stampi, 1992; Tassi & Muzet, 2000). Regarding neural activity, electroencephalography (EEG) studies have demonstrated that the minutes immediately following awakening are characterized by distinct oscillatory changes: specifically, an increase in low-frequency power (delta and theta bands) alongside a reduction in high-frequency power (alpha and beta bands) (Ferrara et al., 2006; Marzano et al., 2011; Vallat et al., 2019).

Many studies on SI have measured behavioral and neural indices at relatively long-time intervals. For example, a recent multimodal imaging study by Vallat et al. (2019) assessed behavior and brain activity (EEG and fMRI) at a pre-sleep baseline and at 5 and 25 minutes after awakening, reporting differences between the SI state (5 minutes) and the recovering state (25 minutes). Some other studies have also measured data at approximately 5-minute intervals even at the finest resolution (e.g., Hayashi et al., 2003; Hilditch et al., 2016, 2023; Hofer-Tinguely et al., 2005; Jewett et al., 1999). The general and coarse-grained trajectory of recovery has begun to be elucidated separately at the regional level (from anterior to posterior: Marzano et al., 2011) and at the frequency level (progressing from theta to alpha, and finally beta: Hilditch et al., 2023). However, it remains largely unknown how neural activity associated with SI reflects combinations of these dimensions—namely, the extent to which specific frequency bands are tied to particular brain regions—and how these neural dynamics temporally evolve across the overall recovery process. Furthermore, the relationship between these neural dynamics and the timing of behavioral recovery, as well as their dependence on the sleep stage immediately prior to awakening, remains unclear.

The present study aims to clarify the temporal dynamics of neural recovery during SI. Specifically, we seek to identify region- and frequency-specific neural activities associated with recovery and to examine their relationship with the timing of behavioral recovery. To this end, we employ a high-temporal-resolution, trial-level approach with 20-second sampling intervals to capture rapid changes in behavioral performance and EEG activity immediately after awakening (Figure 1). We also test whether the depth of pre-awakening sleep (shallow vs. deep NREM) modulates these behavioral and neural dynamics. We predict a temporal sequence in which slow-frequency activity in anterior regions dissipates first, followed by the restoration of faster activity in posterior regions. Furthermore, we hypothesize that these SI-related effects are more pronounced in participants awakened from deep NREM sleep than in those awakened from shallow NREM sleep.

**Figure 1.**
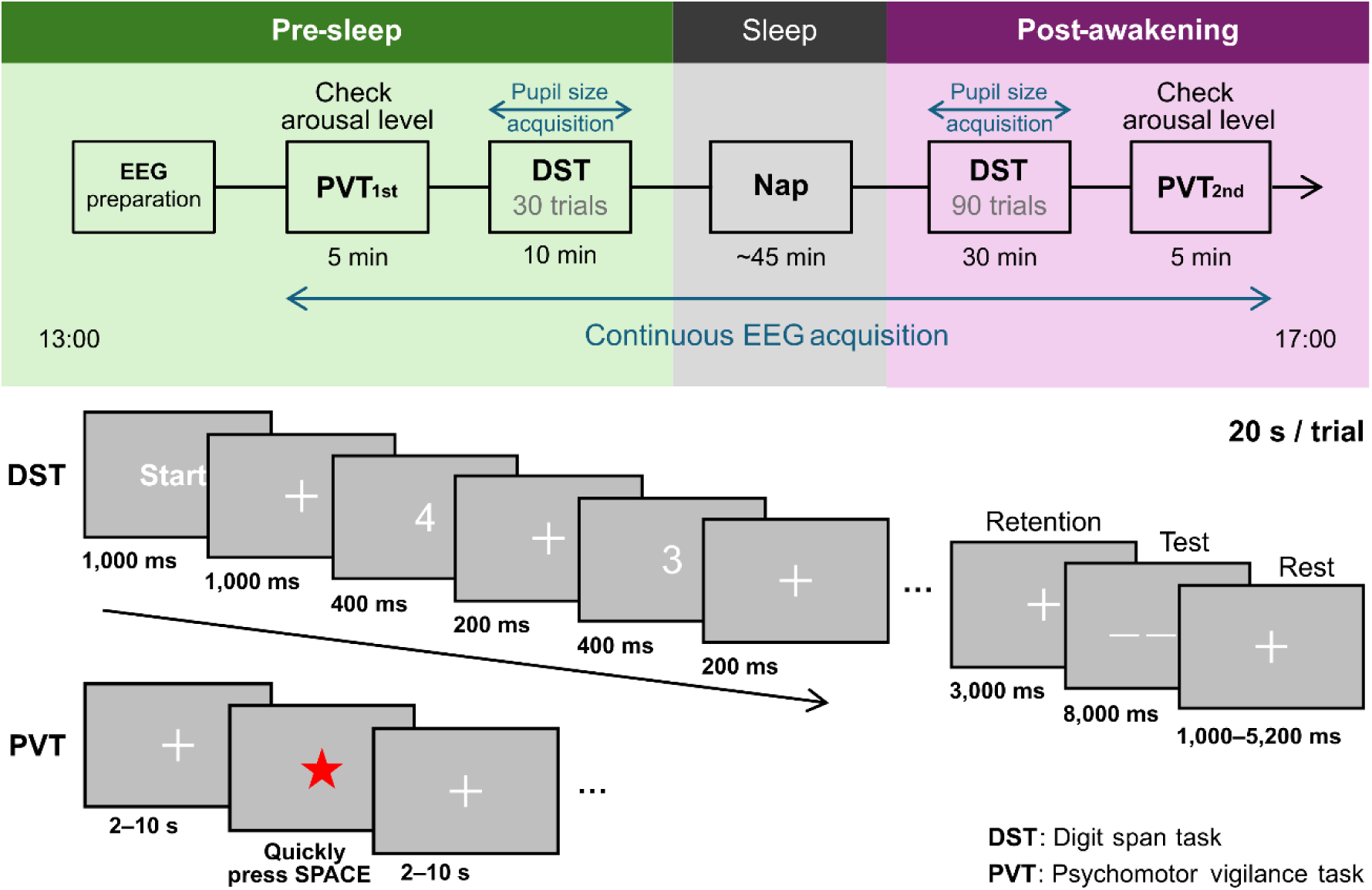
Experimental procedure and task design. Top panel: timeline of the experimental session. DST: Participants viewed a sequence of randomly ordered digits and were required to repeat the sequence in reverse order. Each trial included a 3-second retention interval followed by an 8-second test period. Using a staircase procedure, the sequence length increased by one digit after a correct response and decreased by one after an incorrect response. Each trial lasted exactly 20 s, with the duration of the rest adjusted to compensate for sequence length (1,000–5,200 ms). Pupil size was recorded during the DST in both the pre-sleep and post-awakening periods. PVT: While fixating on a central cross, participants responded to a target (red star) that appeared at jittered inter-stimulus intervals of 2–10 s. Participants were instructed to respond as quickly as possible.

## Methods

### Participants

Sixty-four healthy adults (mean age = 20.14 ± 2.02 years, 16 females) participated in the study. Inclusion criteria were: (i) no history of neurological or psychiatric disorders, (ii) normal or corrected-to-normal vision, and (iii) regular sleep–wake schedule. Participants were excluded in a two-step procedure. First, individuals who were awakened from REM sleep or who were awake at the scheduled awakening time were excluded to restrict analyses to NREM awakenings. Second, additional participants were excluded due to poor EEG data quality (e.g., excessive noise or movement-related artifacts). After these exclusions, the final analyzed sample consisted of 44 participants. The study protocol was approved by the Research Ethics Committee of Kochi University of Technology, and all participants provided written informed consent. All procedures were conducted in accordance with the Declaration of Helsinki.

### Experimental design & procedure

Experimental sessions were conducted in the afternoon between 13:00 and 17:00. Each session consisted of several steps: EEG preparation, baseline behavioral assessments before sleep, a nap, and post-awakening behavioral assessments (Figure 1). EEG recordings were continuously acquired from the pre-sleep period through the post-awakening period.

#### Pre-sleep period

Participants first underwent EEG preparation. Baseline arousal was quantified using a 5-minute psychomotor vigilance task (PVT). In each PVT trial, a fixation cross presented for a variable duration ranging from 2 to 10 s. Upon the appearance of a red star (target stimulus), participants were required to respond as rapidly as possible by pressing a space key. After target onset, either a key press or a 2-s timeout advanced the task to the next trial. The task consisted of 45 trials and lasted approximately 5 minutes.

Before performance assessment, participants reported their subjective sleepiness on a 5-point scale (1 = very sleepy, 5 = not sleepy at all). We employed a backward digit span task (DST), which engages multiple cognitive processes (St Clair-Thompson & Allen, 2013), to robustly capture the effects of SI. Baseline working-memory performance was assessed using a 30-trial DST in a darkened room. To ensure a constant session duration across a fixed number of trials, each trial was fixed at 20 s, resulting in a 10-minute session (30 trials). The DST began with a sequence of three digits; if a participant responded correctly, the digit span increased by one in the subsequent trial, whereas an incorrect response or failure to respond within the allotted time resulted in a decrease of one digit. Each 20-s trial consisted of a fixation period (2.0 s), digit sequence presentation (0.6 s per digit), a retention interval (3.0 s), a test period (8.0 s), and a rest period (1.0–5.2 s). To maintain a constant trial duration of 20 s, the final rest period was adjusted based on the number of digits presented. For instance, a three-digit trial included a 1.8-s display and a 5.2-s rest, whereas a five-digit trial involved a 3.0-s display and a 4.0-s rest. The number of digits presented on each trial was adjusted according to the participant’s performance. Additionally, mean pupil size recorded during the pre-sleep DST served as a baseline physiological marker.

#### Nap period

Following the pre-sleep baseline assessments, participants underwent a monitored nap session in a controlled sleep environment (Figure 2). After a nap period of approximately 45 minutes, participants were abruptly awakened by experimenters and seated as quickly as possible in a chair positioned in front of a computer located immediately beside the bed.

**Figure 2.**
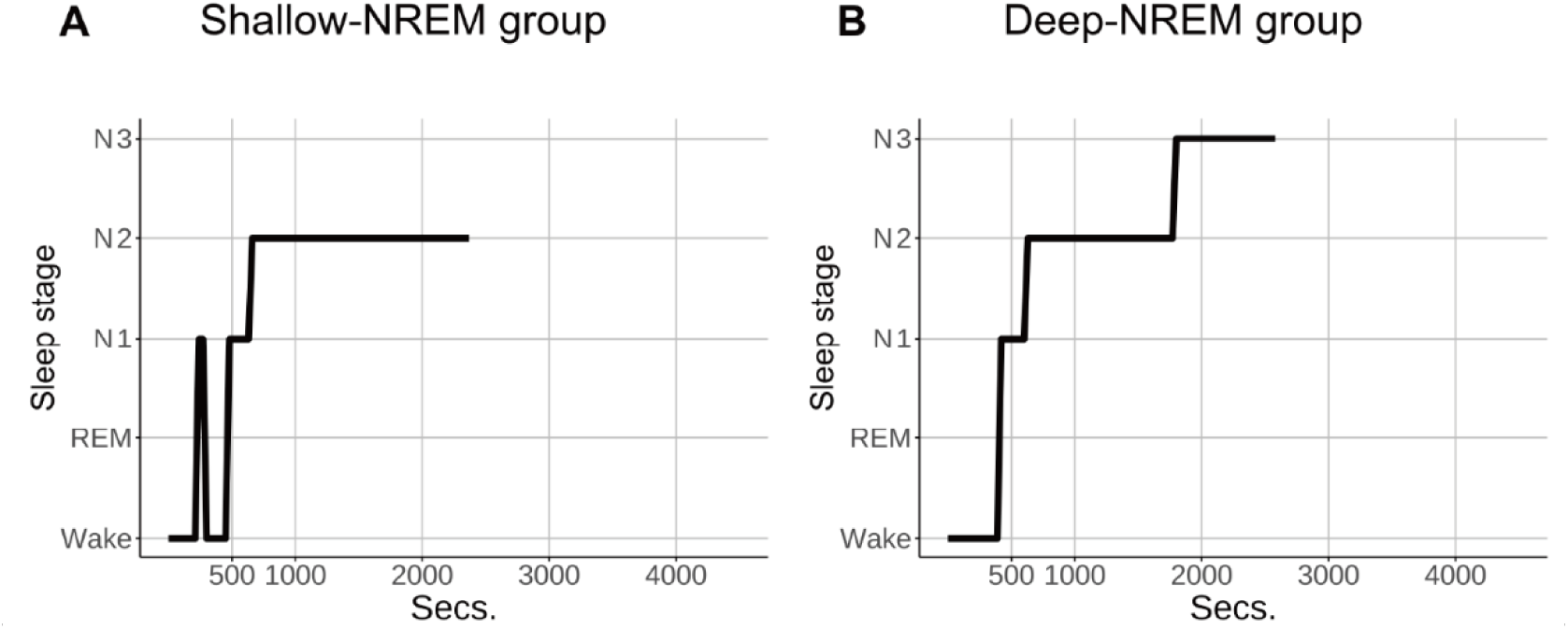
Representative hypnograms for the shallow- and deep-NREM groups. (A) Example hypnogram from a participant in the shallow-NREM group, showing a nap episode ended by N2 sleep. (B) Example hypnogram from a participant in the deep-NREM group, illustrating consolidated progression from wakefulness through N1 and N2 into sustained N3 sleep prior to awakening.

#### Post-awakening period

To quantify SI impairment, working-memory performance was assessed throughout the 30-minute period immediately following awakening. Specifically, participants performed an extended DST consisting of 90 trials in a darkened room. Pupil size was continuously recorded during this DST. After completing this task, participants rated their current subjective sleepiness on a 5-point scale (1 = very sleepy, 5 = not sleepy at all). In addition, participants retrospectively reported the level of subjective sleepiness they experienced immediately after awakening on a 5-point scale (1 = very sleepy, 5 = not sleepy at all), as well as subjective sleep quality on a 5-point scale (1 = slept very well, 5 = did not sleep at all). Finally, arousal level was assessed using a 5-minute PVT, conducted under conditions identical to the baseline assessment.

### Data acquisition and analysis

#### Behavioral data acquisition

The PVT was administered using PsychoPy (https://www.psychopy.org/), whereas the DST was implemented using a custom script written in MATLAB (MathWorks, Natick, MA, USA). Pupil size was continuously recorded at 100 Hz using a Tobii Pro Glasses 3 eye-tracking system (Tobii AB, Sweden).

#### EEG data acquisition

EEG data were recorded using a 32-channel Brain Products system (Brain Products GmbH, Germany) at a sampling rate of 5,000 Hz. Electrodes were positioned according to the international 10–20 system (Klem et al., 1999), with Fpz used as the reference electrode. All electrode impedances were kept below 5 kΩ.

#### Behavioral data analysis

To quantify the time point at which behavioral performance returned to baseline, we defined a return-to-baseline index, Trial_RTB_. This was designated as the first post-awakening trial in which a participant’s performance returned to the pre-sleep baseline level. Baseline performance was operationalized as the maximum digit span achieved during the pre-sleep DST session. Thus, Trial_RTB_ serves as a temporal marker indicating the point at which working-memory capacity was successfully restored to the pre-sleep level. Group differences in Trial_RTB_ were evaluated using a Wilcoxon rank-sum test.

Reaction times (RTs) in the DST were calculated based only on the first digit response in each trial. Trial-level RTs in the post-awakening DST were normalized by subtracting the mean RT from the pre-sleep DST. These indices were analyzed using a generalized linear mixed-effects model (GLMM) with a gamma distribution and a log link function. The model included *Group* (shallow- or deep-NREM) and *Trial* (normalized trial number) factors and their interaction as fixed effects, with a random intercept for participants to account for individual differences in baseline RT and the within-subject dependence of observations.

Pupil size data recorded during the post-awakening DST were normalized by subtracting the mean pupil size obtained during the pre-sleep DST. These data were analyzed using a GLMM with gaussian distribution. The model included *Group*, *Trial*, and their interaction as fixed effects, and a random intercept for participants to capture individual differences in baseline pupil size.

RTs on the PVT were analyzed using a mixed-design ANOVA with *Group* as a between-subjects factor and *Period* (pre-sleep or post-awakening) as a within-subjects factor.All behavioral results were corrected for multiple comparisons using the false discovery rate (FDR) procedure.

#### Sleep stage classification

Sleep stages were classified offline using a pre-trained convolutional recurrent neural network model (CRNN; Abou Jaoude et al., 2020), which estimates sleep stages in 30-second epochs. Based on the predominant sleep stage during the 3-min window immediately preceding awakening, participants were assigned to one of two groups. Specifically, participants who spent the majority of this period in stages N1 or N2 were categorized into the shallow-NREM group, whereas those with a predominance of stage N3 were categorized into the deep-NREM group. Participants who were primarily in a state of wakefulness or REM were excluded from further analysis.

#### EEG preprocessing and spectral decomposition

EEG preprocessing and time-frequency analysis were performed using MNE-Python (Version 1.7.1; https://mne.tools/1.7/), an open-source toolbox for MEG/EEG analysis in Python (Gramfort et al., 2013). Continuous EEG data were first bandpass filtered (0.1–30 Hz) and subsequently down-sampled to 250 Hz. Artifacts were removed using a pipeline combining independent component analysis with manual visual inspection. The data were then segmented into 20-second epochs time-locked to the onset of each DST trial, covering pre-sleep baseline and post-awakening recovery periods. Note that, for each trial, a 1-second pre-stimulus fixation period was used as baseline for normalization. Trials still contaminated by residual artifacts were excluded from further analysis.

Spectral decomposition was conducted using a continuous wavelet transform with complex Morlet wavelets to estimate time-frequency power in the 1–30 Hz range. To minimize inter-trial and inter-channel variability, EEG signals were baseline-normalized using single-trial z-score normalization relative to the mean and standard deviation of the 1-second pre-stimulus fixation period. Subsequently, mean EEG power was quantified for five frequency bands: delta (1–4 Hz), theta (4–8 Hz), alpha (8–12 Hz), beta1 (12–16 Hz), and beta2 (16–30 Hz). To verify that the current data replicated established EEG signatures of SI, band-limited power was averaged across trials and electrodes, and then compared between pre-sleep and post-awakening (0–10 min after awakening) periods using a paired t-test across participants (collapsed across sleep groups).

#### EEG regional analysis and ERSP

For regional analysis, channels were grouped into four topographical regions: frontal (Fp1, Fp2, F3, F4, FC1, FC2, Fz, FC5, FC6, F7, F8), parietal (C3, C4, Cz, CP1, CP2, CP5, CP6, P3, P4, P7, P8, Pz), temporal (T7, T8, TP9, TP10), and occipital (POz, O1, O2, Oz). Event-related spectral perturbation (ERSP) was calculated for each trial as the change in time-frequency power relative to the 1-second pre-stimulus fixation baseline, in line with standard ERSP procedures (Grandchamp & Delorme, 2011). The resultant ERSP was averaged across the temporal dimension to obtain a frequency vector for each trial. These vectors were then averaged within each region and concatenated trial-by-trial to visualize the frequency × trial dynamics of the recovery process.

#### EEG statistical analysis and recovery trial index

To identify significant changes in EEG power after awakening, a cluster-based permutation test was conducted for each region (Maris & Oostenveld, 2007). Briefly, for each frequency–trial bin, t-values were computed comparing post-awakening power to pre-sleep baseline power. Clusters were defined as contiguous bins exceeding a cluster-forming threshold of *P* < 0.01 (two-tailed). Cluster significance was then assessed by comparing summed t-values within each cluster against the null distribution generated from 5,000 permutations (*P* < 0.05).

To further characterize the temporal dynamics of these changes, we calculated a recovery trial index. For each significant frequency × trial cluster, we first averaged the post-awakening power within the frequency bands trial-by-trial. The recovery trial was defined as the first trial in which the mean power no longer differed significantly from the average power across all 30 pre-sleep baseline trials (*P* ≥ 0.05, paired t-test, FDR-corrected) and remained non-significant for all subsequent trials, ensuring the identification of sustained functional restoration.

## Results

### Sleep characteristics and subjective measures

The temporal organization of sleep stages was comparable between shallow- and deep-NREM groups. Participants in the shallow-NREM group showed a mean total sleep time of 44.1 ± 9.0 min, which did not differ significantly from that in the deep-NREM group (40.9 ± 6.4 min, *t* = 1.4, *P* = 0.168). Figure 2 shows representative hypnograms from the shallow- and deep-NREM groups.

To confirm participants’ subjective state, we examined self-reported sleepiness and sleep quality. Participants rated their sleepiness at three time points: before the pre-sleep DST, immediately after awakening, and after the post-awakening DST. No significant group differences were observed at any time point (shallow NREM: before pre-sleep DST, M = 2.32, SD = 0.95; immediately post awakening, M = 2.21, SD = 1.03; after post-awakening DST, M = 3.57, SD = 1.08; deep NREM: before pre-sleep DST, M = 2.82, SD = 1.05; immediately post awakening, M = 2.73, SD = 1.32; after post-awakening DST, M = 3.39, SD = 1.34; Welch’s tests, all ps > 0.1, Holm-corrected). Similarly, subjective sleep-quality ratings did not differ between groups (shallow NREM: M = 2.14, SD = 0.91; deep NREM: M = 1.96, SD = 1.02; Welch’s test, p > 0.1).

Taken together, these results demonstrate that the groups were well-matched in overall sleep characteristics, with the primary difference residing in the sleep stage at the point of awakening.

### Behavioral results

Behavioral recovery following awakening varied as a function of sleep depth. In the DST, participants awakened from deep NREM sleep required a significantly greater number of trials to return to their pre-sleep digit-span performance (Trial_RTB_) compared to those awakened from shallow-NREM sleep (Wilcoxon rank-sum test, *W* = 298.5, *P* = 0.0194; Figure 3). This finding indicates that working-memory recovery is substantially protracted following deep NREM sleep. Trial-level RTs decreased significantly over time (GLMM, main effect of *Trial*, *P* < 0.001), but neither the main effect of *Group* (shallow- or deep-NREM) nor the *Group* × *Trial* interaction reached significance (*P* > 0.1), indicating a similar rate of RT improvement between groups (Figure S1). Pupil size increased throughout the post-awakening period (GLMM, main effect of *Trial*, *P* < 0.001). A significant *Group* × *Trial* interaction (*P* = 0.002) revealed a steeper rate of pupil dilation in the shallow-NREM group, consistent with a more rapid restoration of physiological arousal compared to the deep-NREM group (Figure S2).

**Figure 3.**
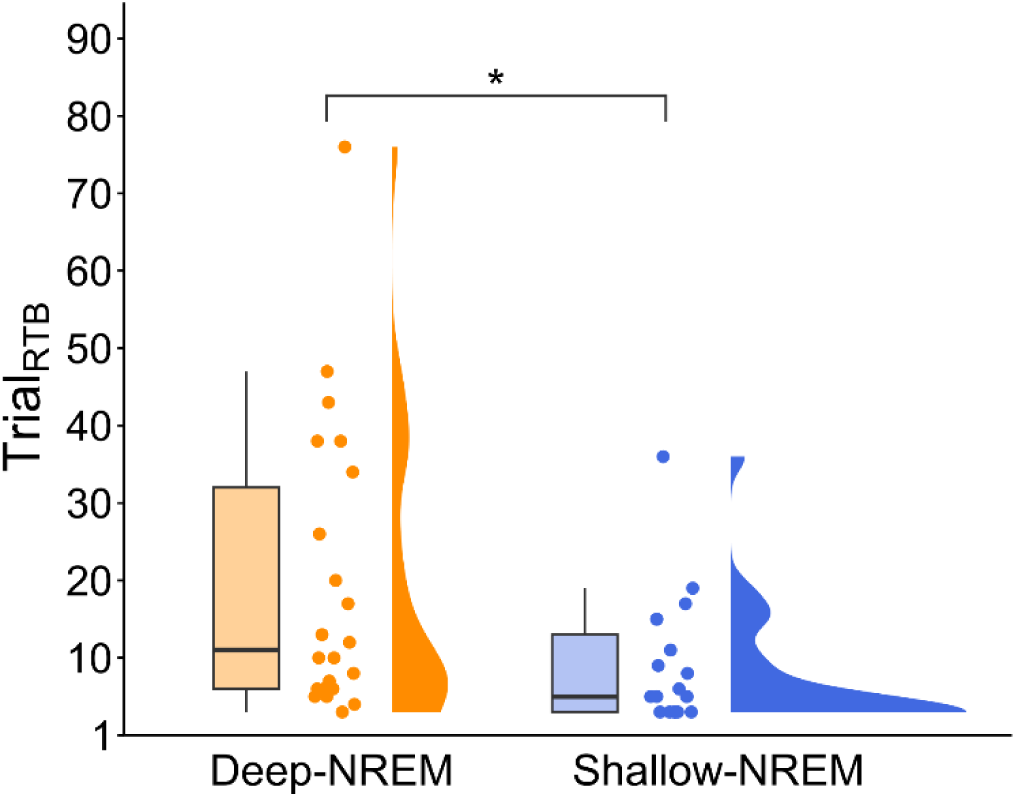
Trial_RTB_ for deep- and shallow-NREM groups. The number of trials required for working-memory performance to return to the pre-sleep level (i.e., Trial_RTB_) is separately shown for deep- and shallow-NREM groups. The box plot represents the median value (thick line), the interquartile range (box), and the minimum and maximum values (whiskers). Raw data points and smoothed distribution are also shown. *, *P* < 0.05.

For RTs in the PVT, a mixed-design ANOVA revealed no significant main effect of *Group*, *Period* (pre-sleep or post-awakening), or *Group* × *Period* interaction (*P* > 0.1) (Figure S3).

### Whole-scalp EEG spectral characteristics following awakening

We first sought to replicate the canonical spectral patterns of SI to confirm the validity of our recordings. Statistical comparisons between pre-sleep and post-awakening power revealed a pattern of frequency-specific modulations (Figure 4).

**Figure 4.**
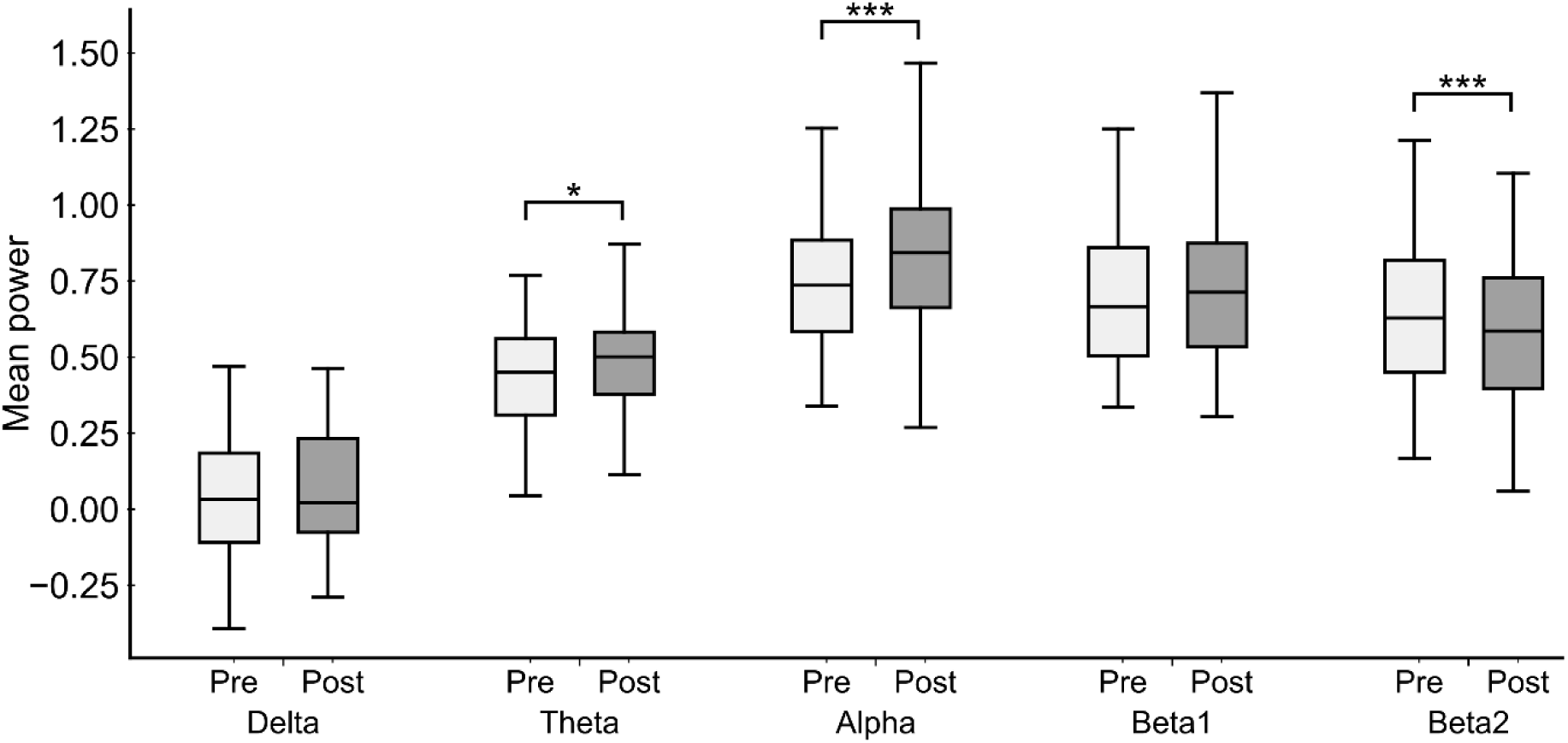
Whole-scalp EEG power. Boxplots illustrate EEG power during the pre-sleep and post-awakening (0–10 min from the start of DST) periods for five frequency bands, averaged across all electrodes. *, *P* < 0.05. ***, *P* < 0.001.

Delta (1–4 Hz) power did not change significantly (*t* = 0.87, *P* = 0.39, FDR-corrected), nor did beta1 (12–16 Hz) power (*t* = 1.86, *P* = 0.087, FDR-corrected). In contrast, we observed a significant elevation in theta (4–8 Hz; *t* = 2.34, *P* = 0.040, FDR-corrected) and alpha power (8–12 Hz; *t* = 4.91, *P* < 0.001, FDR-corrected), alongside a marked reduction in beta2 power (16–30 Hz; *t* = 4.27, *P* < 0.001, FDR-corrected). These findings—specifically the simultaneous increase in slower oscillations and suppression of higher-frequency beta activity—are in full alignment with established models of the awakening process (Ferrara et al., 2006; Gorgoni et al., 2015, 2019; Landolt et al., 2019; Marzano et al., 2011; Tassi & Muzet, 2000). This robust replication of the characteristic SI profile provides a reliable foundation for our subsequent trial-level analyses.

### Sleep-depth dependent spatiotemporal dynamics of EEG power following awakening

Next, to determine whether recovery trajectories differ according to prior sleep depth, we stratified the analysis of post-awakening EEG dynamics into deep-NREM (N3) and shallow-NREM (N1/N2) groups. This approach facilitated a more fine-grained evaluation of how sleep depth modulates regional recovery processes across the cortex.

In the deep-NREM group, we observed distinct region- and frequency-specific modulations of post-awakening EEG activity. In the frontal region, delta-to-theta band power (approximately 2–7 Hz) exhibited a significant reduction immediately after awakening (*P* < 0.05, cluster-corrected; Figure 5A). This suppression was transient, localized to the initial trials (∼1 min), and gradually returned to baseline thereafter. In the parietal region, a significant decrease in high-frequency beta-band activity (approximately 15–30 Hz) coincided with a sustained increase mainly in alpha activity (approximately 5–14 Hz) (*P* < 0.05, cluster-corrected; Figure 5B). These effects were most pronounced during the first 20–25 trials (at approximately 7.5–8.5 min) and diminished progressively over time. Similarly, the temporal region exhibited a marked elevation in theta-to-alpha band power (approximately 2–13 Hz, dominated by the alpha band) during the early post-awakening phase [trials 10–25 (approximately 3–8 min); *P* < 0.05, cluster-corrected], which subsequently declined as the session progressed (Figure 5C). Finally, the occipital region demonstrated a robust increase in alpha activity during the initial 20 trials (∼7 min; *P* < 0.05, cluster-corrected; Figure 5D).

**Figure 5.**
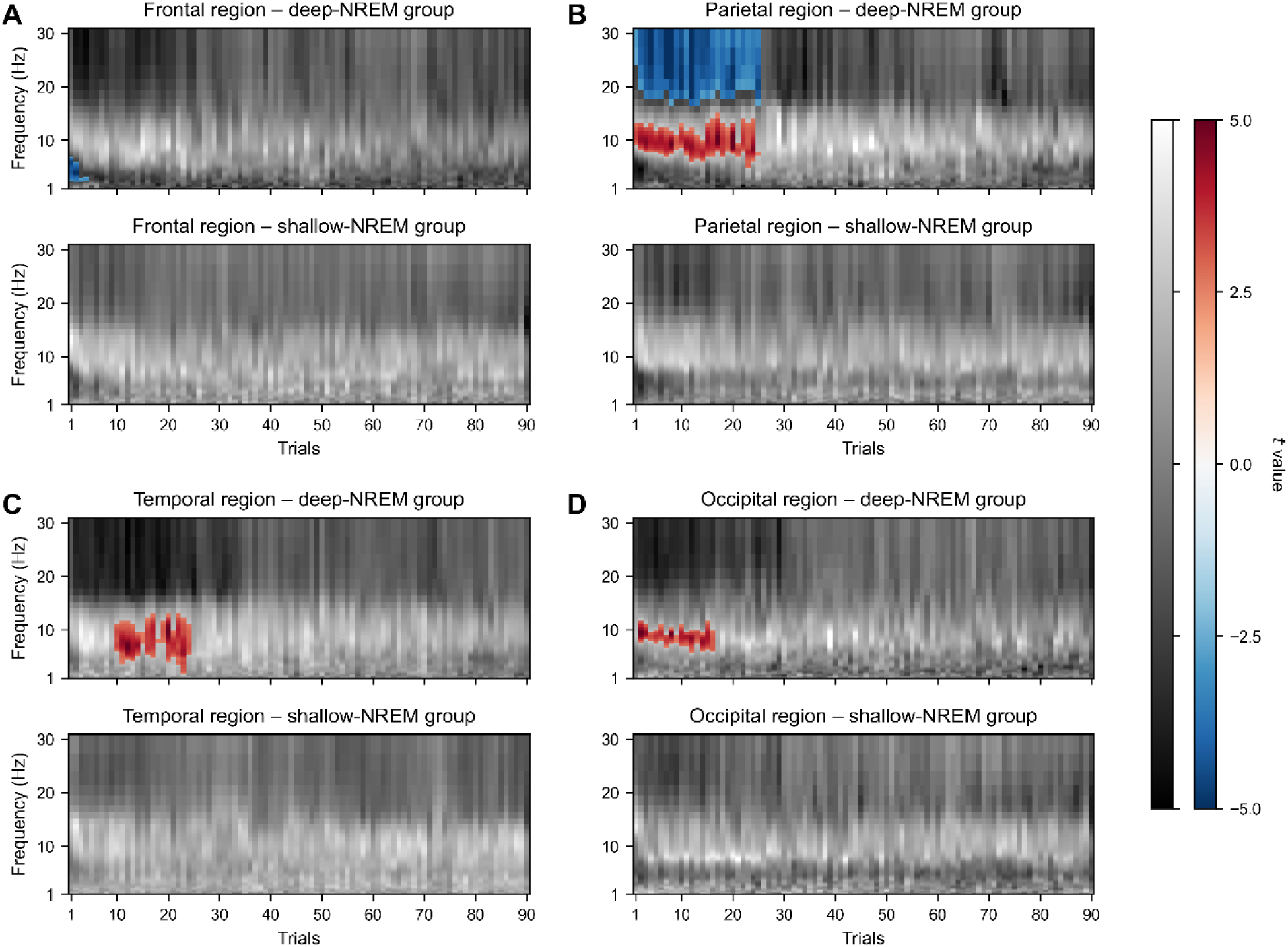
Spatio-temporal dynamics of post-awakening EEG power across cortical regions. The time-frequency maps (t-values against pre-sleep baseline) are shown for the frontal (A), parietal (B), temporal (C), and occipital (D) regions. For each region, results are presented separately for the deep-NREM (top) and shallow-NREM (bottom) groups. Significant clusters identified via cluster-based permutation test (cluster-forming threshold of *P* < 0.01; cluster-corrected threshold of *P* < 0.05) are indicated by colored areas. Red and blue denote power increases and decreases relative to pre-sleep baseline, respectively. Black-and-white shading represents t-values for non-significant areas. Data were averaged across electrodes within each brain region and normalized as described in the Methods section. Clusters spanning at least three consecutive trials are shown.

In contrast, the shallow-NREM group exhibited no significant EEG clusters across any cortical region or frequency band (Figure 5A–D), suggesting that neural dynamics following awakening from shallow-NREM sleep are substantially less disrupted than those following deep-NREM sleep. This stability in the spectral profile is consistent with our behavioral finding of attenuated SI observed after shallower sleep.

By employing the recovery trial index, we confirmed that EEG recovery did not occur uniformly across cortical regions and bands (Figure 6; for details, see the Methods section). Frontal delta/theta activity returned to baseline by the 4^th^ trial (1.33 min from the onset of the DST), representing the earliest normalization among all examined regions. In contrast, posterior and lateral regions exhibited more extensive lags: occipital alpha power recovered by the 16^th^ trial (5.33 min), whereas temporal alpha and parietal alpha/beta2 activity remained significantly deviated from baseline until the 23^rd^ and 25^th^ trials (7.67 and 8.33 min), respectively. These recovery indices reveal a graded spatiotemporal pattern in which frontal slow activity normalizes first, followed by mid-frequency activity in occipital regions, whereas mid and higher-frequency activity in temporal and parietal regions exhibits the most protracted recovery trajectories. Behavioral recovery in the deep-NREM group was temporally situated between occipital and parieto-temporal recovery.

**Figure 6.**
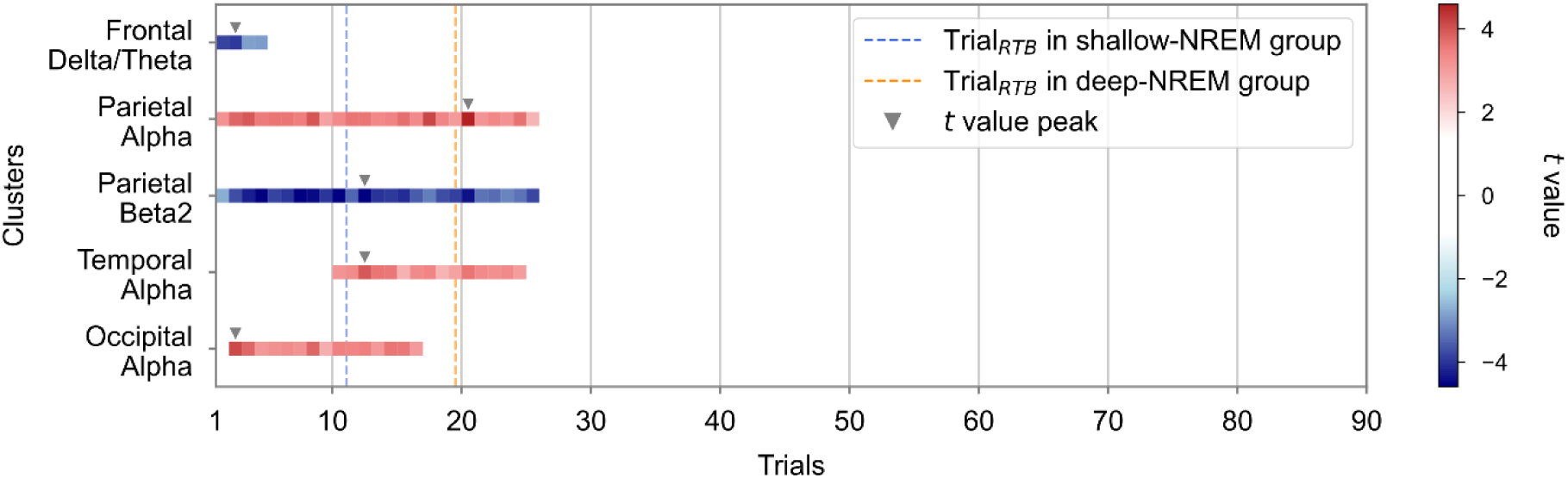
Temporal dynamics of region-specific spectral power in the deep-NREM group. Significant clusters (identified in Figure 5) were averaged across each frequency band, with the resulting test statistics (relative to the pre-sleep baseline) plotted on a trial-by-trial basis. Red and blue bins denote significantly increased and decreased power relative to baseline, respectively; thus, the cessation of a continuous sequence of colored bins marks the recovery trial. Dashed vertical lines indicate the trial at which working-memory performance returned to baseline in DST (i.e., Trial_RTB_) for each group (blue: shallow-NREM; orange: deep-NREM). Triangle denotes the trial with the peak t-value within each cluster.

## Discussion

We characterized the recovery process from SI at high temporal resolution using fine-grained behavioral measurements and EEG analyses. By categorizing the pre-awakening NREM sleep stage based on depth, we found that individuals who awakened from deep NREM sleep exhibited more pronounced behavioral and neural profiles. At the behavioral level, SI exerted a clear inhibitory effect on working-memory performance, with a significantly longer duration of impairment in the deep-NREM group. This pattern was also reflected in pupil size data. At the neural level, the deep-NREM group showed an immediate post-awakening decrease in parietal beta2 activity, a reduction in frontal slow activity, and an increase in alpha activity across parietal, temporal, and occipital regions. These effects were not observed in the shallow-NREM group. In the deep-NREM group, behavioral recovery occurred approximately 7 minutes after awakening, following the restoration of frontal slow activity and occipital alpha power to baseline levels (Figure 6). In contrast, activities in parietal and temporal regions returned to baseline only after behavioral recovery had been achieved. Together, this tight temporal coupling between neural and behavioral changes suggests a coordinated functional process underlying human recovery from SI, providing a strong foundation for elucidating the neural mechanisms.

### Parietal beta suppression and cognitive “unreadiness”

A hallmark of the neural profile in the deep-NREM group was the significant reduction of beta power in the parietal region during the early post-awakening period. Given beta activity was considered to reflect cortical activation (Steriade et al., 1990) and arousal levels (Cardenas et al., 1997), beta-band suppression during SI has been widely interpreted as a signature of attenuated cortical arousal and a lack of readiness for cognitive processing (Ferrara et al., 2006; Gorgoni et al., 2015; Marzano et al., 2011; Vallat et al., 2019). Our results extend this view by demonstrating that beta suppression is particularly pronounced over the parietal cortex. Given the parietal lobe’s central role in attentional selection, working-memory updating, and the integration of task-relevant information (Corbetta & Shulman, 2002; Owen et al., 2005; Wager & Smith, 2003) —functions directly engaged by the DST paradigm—this localized suppression may reflect a specific state of “unreadiness” within posterior attentional hubs. This interpretation is consistent with the protracted behavioral recovery observed in the deep-NREM group, suggesting that parietal beta dynamics may constitute a key neural correlate of delayed restoration of task-related cognitive capacity following deep NREM sleep.

### Frontal delta/theta dissipation and rapid mobilization

Although we observed a slight increase in delta/theta activity at the scalp-averaged level, which was consistent with previous studies (Ferrara et al., 2006; Hilditch et al., 2023; Tomzig et al., 2025), a distinctive feature in the deep-NREM group was the immediate reduction of frontal delta/theta activity. This aligns with reports of localized frontal low-frequency power decreases (Marzano et al., 2011; Radzi et al., 2018). We propose that this reduced frontal slow power reflects an early arousal-related “dissipation” of sleep-like activity, marking the onset of wake-like frontal dynamics. This view is supported by studies highlighting the critical role of frontal vigilance networks upon awakening (Balkin et al., 2002; Chen et al., 2020; Wang et al., 2024). Alternatively, the immediate demands of the DST may have elicited rapid prefrontal engagement, thereby suppressing slow power through task-induced mobilization. Our results suggest that frontal slow power suppression serves as a neural signature of rapid, potentially externally driven, cognitive mobilization during the initial stage of awakening.

### Posterior alpha elevation and topographical dissociation

Within parietal, occipital, and temporal regions, the deep-NREM group exhibited a persistent increase in alpha-band power. Alpha increase has been associated with reduced vigilance, attentional disengagement, and increased subjective sleepiness (Foxe & Snyder, 2011; Klimesch, 1999; Mathewson et al., 2011; Radzi et al., 2018; Santamaria & Chiappa, 1987). Thus, the sustained alpha elevation in these cortices likely reflects a protracted state of reduced perceptual and attentional readiness that outlasts the initial behavioral impairments, consistent with SI work showing persistence of sleep-like EEG features after awakening, particularly in posterior regions (Ferrara et al., 2006). Conversely, the absence of alpha increase in frontal regions, coupled with the early reduction of frontal slow activity, suggests that prefrontal networks, involved in executive control, may recover more rapidly or be preferentially engaged by task demands (e.g., prefrontal activity supporting overcoming sleepiness under task load; Honma et al., 2010). This topographical dissociation implies that while the executive “engine” of the brain re-engages quickly, posterior sensory and attentional hubs remain in a lingering state of sleep-like hypo-arousal.

### Hierarchical spatiotemporal cortical recovery dependent on sleep depth

While prior research has independently elucidated recovery trajectories at the frequency (Hilditch et al., 2023) or regional level (Marzano et al., 2011), our results reveal that cortical recovery follows a distinct, integrated spatiotemporal hierarchy. This process involves the rapid normalization of frontal low-frequency activity contrasting with the substantially protracted recovery of high-frequency activity in posterior regions (occipital, temporal, and parietal). This “frontal-first, posterior-last” gradient aligns with evidence of asynchronous cortical arousal (Stephan et al., 2025) and likely reflects the structural organization of arousal systems. Crucially, this graded pattern of extensive cortical reorganization was exclusive to awakenings from deep NREM sleep (N3), whereas awakenings from shallow NREM sleep (N1/N2) exhibited minimal spectral perturbations. This dissociation suggests that pre-awakening sleep depth is the critical determinant dictating the magnitude of neural reorganization required to achieve full wakefulness.

### Limitations and future directions

Several limitations warrant consideration. First, we examined SI following daytime naps. Although behavioral impairments are reportedly comparable to those following nocturnal awakenings (Achermann et al., 1995), it remains unclear whether the underlying neural dynamics are identical. Second, awakenings from REM sleep were excluded. Given that REM-related SI can be severe and involve distinct emotional reactivity (Marzano et al., 2011; Occhionero et al., 2021), future research should directly compare recovery trajectories between REM and NREM awakenings using high-temporal-resolution paradigms.

The precise functional roles of the observed spatiotemporal spectral shifts remain to be elucidated. These dynamics may trigger subsequent recovery across distributed cortical networks (Deco et al., 2019; Hilditch et al., 2023). To map the large-scale network reconfigurations during the sleep-to-wake transition, future studies employing high-spatial-resolution techniques, such as fMRI or simultaneous EEG–fMRI, are required.

## Conclusion

This study demonstrates that awakenings from deep NREM sleep led to the protracted recovery of working-memory performance and pronounced, region- and frequency-specific EEG deviations. The identified patterns—parietal beta suppression, sustained posterior alpha enhancement, and relatively rapid recovery of frontal slow activity—reveal a frequency-specific and regionally asynchronous transition from sleep to wakefulness. These findings highlight the critical importance of specific timescales and sleep depth to fully elucidate the neural mechanisms underlying SI.

## Data availability

All custom Python codes as well as multimodal data are available at https://github.com/masaki-takeda/slp.

## Author contributions

D.S. and M.T. conceived the project. C.H. and D.S. collected data. C.H. and D.S. analyzed data. C.H., D.S., K.N. and M.T. wrote the manuscript.

## Acknowledgments

We would like to thank Ms. Maoko Yamanaka for her administrative assistance. This study was supported by KAKENHI from Japan Society for the Promotion of Science (20H00521 and 21K18267 to M.T., 23K17174 and 24KJ0192 to D.S.), Adaptable and Seamless Technology Transfer Program through Target-driven R&D (A-STEP) from Japan Science and Technology Agency (JST) Grant Number JPMJTR25UE to M.T., a grant from Takeda Science Foundation to M.T., and a grant from Uehara Memorial Foundation to M.T.

