## Supporting Information for "Asynchronous cortical recovery from sleep inertia: region- and frequency-specific dynamics following deep NREM sleep"

This file includes:

Figure S1 to S3

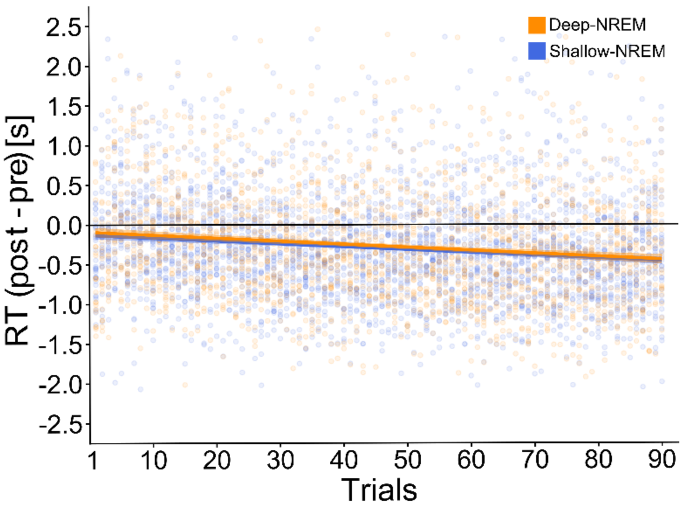


**Figure S1. Reaction time (RT) in the deep-NREM and shallow-NREM groups during the digit span task (DST).** RT for the first response in each post-awakening working-memory trial is shown. To account for individual differences, each RT was normalized by subtracting the participant's grand mean RT in the pre-sleep session. Each dot represents a single response from one participant. Orange and blue lines represent the deep-NREM and shallow-NREM groups, respectively. Error bands denote ± SEM. RT significantly decreased across trials ($P < 0.001$) in both the deep-sleep and shallow-sleep groups. No significant main effect of *Group* was observed ($P = 0.579$), nor a *Group* × *Trial* interaction ($P = 0.618$).

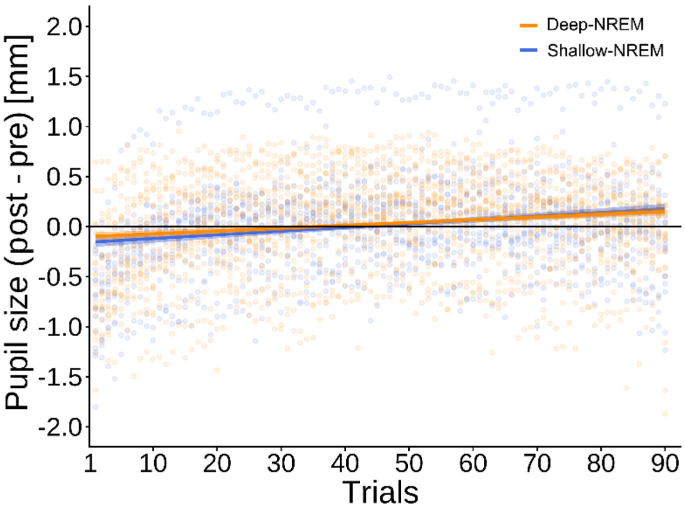


**Figure S2. Mean pupil size across trials in the deep-NREM and shallow-NREM groups.** The average pupil size for each trial is shown. To account for individual differences, pupil size was normalized by subtracting each participant's grand mean pupil size from the pre-sleep session. Each dot represents a single response from one participant. The orange and blue lines represent the deep-NREM and shallow-NREM groups, respectively. Error bands denote ± SEM. The significant *Group* × *Trial* interaction (*P* = 0.002) indicates a steeper time-related increase in the shallow-NREM group compared with the deep-NREM group.

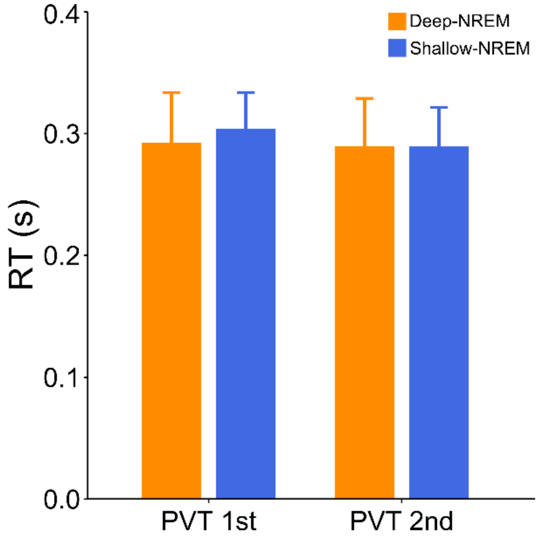


**Figure S3. RT of deep-NREM and shallow-NREM groups during two PVT sessions.** The vigilance of the participants was assessed using a modified version of the psychomotor vigilance task (PVT) at the start and the end of the experiment. There was no significant effect of period or sleep stage on the reaction time, confirming no significant difference in baseline vigilance.
